# Dysregulated Pulmonary Inflammatory Responses Exacerbate the Outcome of Secondary Aspergillosis Following Influenza

**DOI:** 10.1101/2023.06.27.546808

**Authors:** Chrono K. Lee, Lorena V. N. Oliveira, Ali Akalin, Charles A. Specht, Diana Lourenco, Christina L. Gomez, Zaida G. Ramirez-Ortiz, Jennifer P. Wang, Stuart M. Levitz

## Abstract

Inhalation of airborne conidia of the ubiquitous fungus *Aspergillus fumigatus* commonly occurs but invasive aspergillosis is rare except in profoundly immunocompromised persons. Severe influenza predisposes patients to invasive pulmonary aspergillosis by mechanisms that are poorly defined. Using a post-influenza aspergillosis model, we found that superinfected mice had 100% mortality when challenged with *A. fumigatus* conidia on days 2 and 5 (early stages) of influenza A virus infection but 100% survival when challenged on days 8 and 14 (late stages). Influenza-infected mice superinfected with *A. fumigatus* had increased levels of the pro-inflammatory cytokines and chemokines IL-6, TNFα, IFNβ, IL-12p70, IL-1α, IL-1β, CXCL1, G-CSF, MIP-1α, MIP-1β, RANTES and MCP-1. Surprisingly, on histopathological analysis, superinfected mice did not have greater lung inflammation compared with mice infected with influenza alone. Mice infected with influenza had dampened neutrophil recruitment to the lungs following subsequent challenge with *A. fumigatus*, but only if the fungal challenge was executed during the early stages of influenza infection. However, influenza infection did not have a major effect on neutrophil phagocytosis and killing of *A. fumigatus* conidia. Moreover, minimal germination of conidia was seen on histopathology even in the superinfected mice. Taken together, our data suggest that the high mortality rate seen in mice during the early stages of influenza-associated pulmonary aspergillosis is multifactorial, with a greater contribution from dysregulated inflammation than microbial growth.

**Importance:** Severe influenza is a risk factor for fatal invasive pulmonary aspergillosis; however, the mechanistic basis for the lethality is unclear. Utilizing an influenza-associated pulmonary aspergillosis (IAPA) model, we found that mice infected with influenza A virus followed by *A. fumigatus* had 100% mortality when superinfected during the early stages of influenza but survived at later stages. While superinfected mice had dysregulated pulmonary inflammatory responses compared to controls, they had neither increased inflammation nor extensive fungal growth. Although influenza-infected mice had dampened neutrophil recruitment to the lungs following subsequent challenge with *A. fumigatus*, influenza did not affect the ability of neutrophils to clear the fungi. Our data suggest that the lethality seen in our model IAPA is multifactorial with dysregulated inflammation being a greater contributor than uncontrollable microbial growth. If confirmed in humans, our findings provide a rationale for clinical studies of adjuvant anti-inflammatory agents in the treatment of IAPA.

## Introduction

*Aspergillus* species are ubiquitous saprophytic fungi that cause a wide range of diseases from allergic to invasive aspergillosis (1, 2). *A. fumigatus* is the most common species of *Aspergillus* found in the environment and human infections. Inhalation of the aerosolized spores (conidia) is the primary route of exposure to *A. fumigatus*: on average, a person is estimated to inhale hundreds of conidia daily (1–3). Despite frequent exposure, *A. fumigatus* is relatively non-pathogenic in immunocompetent hosts. However, in immunocompromised hosts, the conidia can swell and germinate into the invasive hyphal morphotype with the resultant development of life-threatening invasive aspergillosis (2). Worldwide, >200,000 people/year are estimated to develop invasive aspergillosis (4, 5). Risk factors include neutropenia, chronic granulomatous disease, hematological malignancies, solid organ and hematopoietic stem cell transplants, receipt of immunosuppressant treatments, and AIDS (1, 2).

Influenza A virus (IAV) is a respiratory pathogen responsible for seasonal outbreaks, epidemics, and periodic pandemics. In the United States, influenza causes hundreds of thousands of hospitalizations and tens of thousands of deaths annually (6, 7). Influenza can damage the epithelial cell layer and alter immune cell function (8–11), thereby increasing vulnerability to secondary infections including invasive aspergillosis (6, 7, 12–14). In models of influenza-associated pulmonary aspergillosis (IAPA), mice sequentially challenged with IAV and *A. fumigatus* had increased mortality, reduced neutrophil recruitment to the lung (15), defective phagolysosome maturation, and reduced fungal clearance by the phagocytic cells (16). Recently, Sarden *et al.* (17) found influenza infection induces death of B1a cells, compromising natural antibody production and weakening humoral defenses against *A. fumigatus*. Still the cause of lethality in IAPA remains incompletely defined.

In this study, we used a mouse model to explore the mechanistic underpinnings of the high mortality in IAPA. To avoid overwhelming the host, we challenged the mice with a dose of *A*. *fumigatus* lower than used in other models of IAPA. We found the lethality of IAPA is dependent on the stage of influenza infection, with mortality only seen in mice challenged with *A. fumigatus* during the early stages of influenza infection. Although prior influenza infection did not significantly impact the amount of inflammation in the lungs following *A. fumigatus* superinfection, the inflammatory response appeared to be dysregulated as evidenced by elevated levels of pro-inflammatory cytokines and chemokines and suppressed neutrophil recruitment. Interestingly, scant growth of *A. fumigatus* was seen in the lungs of mice regardless of whether they had influenza infection, and the viral load was not affected by fungal challenge. Thus, the cause of the high mortality rate in our model of IAPA appears to be multifactorial with damage due to dysregulated inflammatory responses predominating over fungal growth.

## Results

### Establishment of influenza and *A. fumigatus* mouse models

Severe influenza infection predisposes patients to secondary invasive aspergillosis infections (12, 13). To create a mouse model that mimics IAPA in humans, we first performed dose-response experiments to optimize inocula of influenza A virus (IAV) and *A. fumigatus*. To determine infectious doses for IAV that resulted in the mice exhibiting signs of infection yet surviving, mice were infected with varying inocula of IAV (Fig. 1A and 1B) following which weight change and survival were monitored over time. All the mice succumbed within 9 days post IAV infection (dpii) at doses 1000 and 2500 PFU (Fig. 1A). At the lower doses, nearly 100% of the mice survived the IAV infection. However, for all doses tested, infected mice lost weight by 3-4-dpii with significant weight loss peaking around 8-dpii. By 10-dpii, the mice started to recover with body weight approaching baseline values by 14-dpii except for the group infected with 250 PFU; these mice still had not recovered their body weight by 21-dpii (Fig. 1B and Supplementary Table 1 for complete statistical analysis).

**Figure 1.**
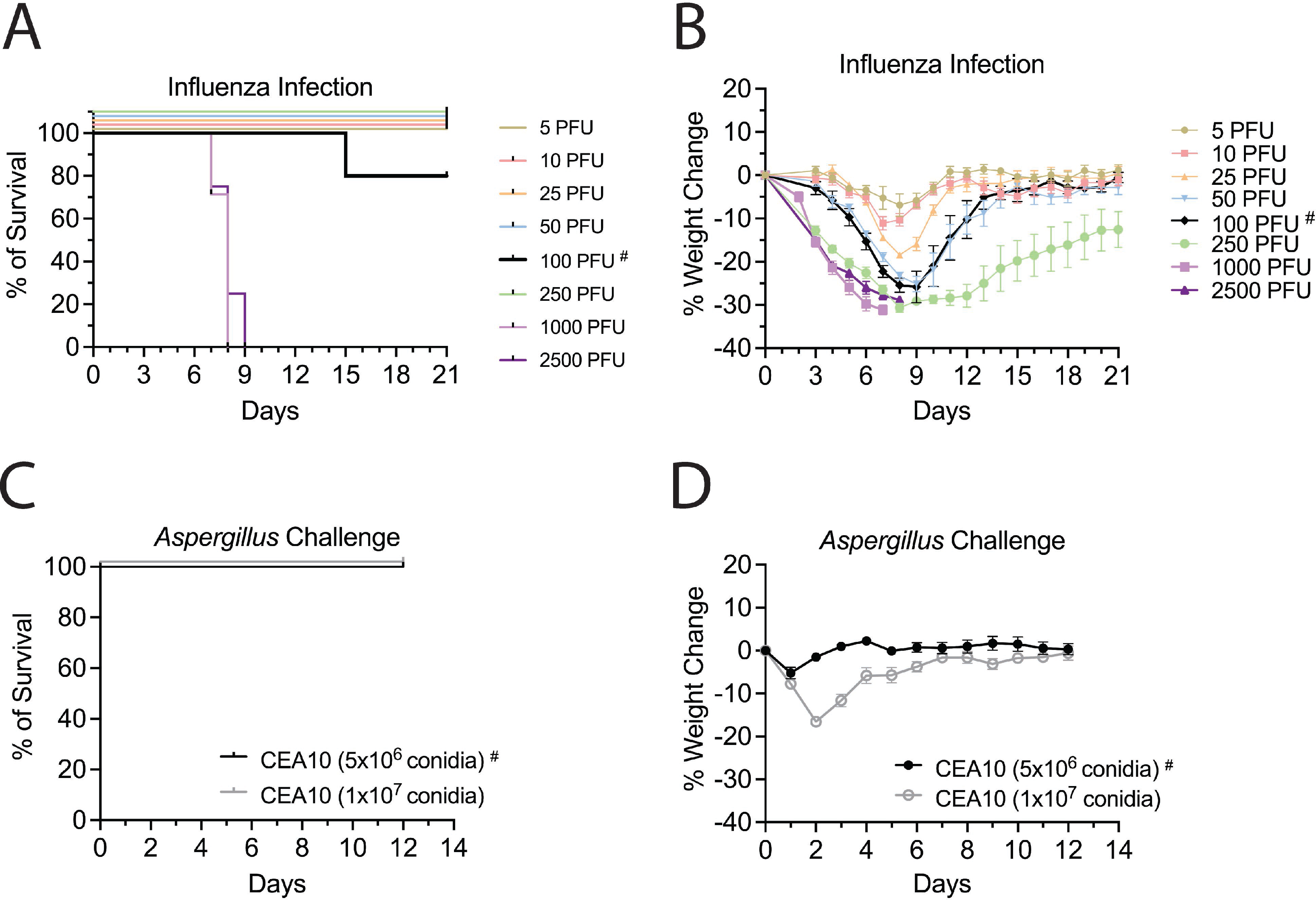
Titration experiments to optimize influenza A virus (IAV) and *Aspergillus fumigatus* (*Af*) infectious inocula. IAV model: Mice were infected with IAV at a range of 5 to 2500 PFU/mouse (C57BL/6) via the I.N. route and monitored daily for survival (A) and body weight change (B). *A. fumigatus* model: Mice were challenged with either 5×10^6^ or 1×10^7^ *A. fumigatus* CEA10 conidia via O.T. route and monitored for survival (C) and body weight change (D). Data represent ≥2 experiments: for IAV model N≥4 mice/group and for *A. fumigatus* model N=12 mice/group. The inocula used for subsequent experiments are indicated by a solid black line and # symbol. Statistical analysis of the weight change curves is shown in Supplementary Table 1.

Previously (18), we showed that wild type (WT) mice had a high tolerance to pulmonary *A. fumigatus* challenge, with near 100% survival even when challenged with 5×10^7^ conidia. However, high inocula of *A. fumigatus* cause significant weight loss and inflammation in mice (16, 17, 19) and may not mimic how hospitalized patients with severe IAV contract secondary aspergillosis. Therefore, we challenged the mice at lower inocula (1×10^7^ and 5×10^6^ conidia) to avoid overwhelming the host. We observed no mortality with these doses (Fig. 1C). However, the mice challenged with 1×10^7^ conidia lost about 15% of their body weight and did not recover their body weight until 7 days post fungal challenge. At the lower dose of 5×10^6^ conidia, the mice lost about 5% of their body weight at day 1 following the challenge but then recovered their weight by day 2 (Fig. 1D and Supplementary Table 1 for complete statistical analysis). Based on these results, we decided to use 100 PFU of IAV and 5×10^6^ *A. fumigatus* conidia as the infectious inocula in subsequent superinfection experiments.

### Mice are more vulnerable to *A. fumigatus* challenge during the early stages of IAV infection compared to the later stages

We next sought to establish a mouse model that could recapitulate the hypersusceptibility of humans with severe influenza to invasive aspergillosis by infecting mice with IAV followed at specified time points by *A. fumigatus*. Thus, on day 0, mice were infected with 100 PFU of IAV. Then, different groups of mice were challenged with 5×10^6^ *A. fumigatus* conidia per mouse at 2-, 5-, 8-, and 14-dpii (Fig. 2A). The mice were monitored daily for percent weight change (Fig. 2B) and survival (Fig. 2C). The mice showed rapid weight loss after *A. fumigatus* challenge at 2- and 5-dpii and eventually succumbed to the superinfection by 5 to 7 days after fungal challenge. There was a delay in recovery of weight for the group that was challenged with *A. fumigatus* at 8-dpii although mortality was not significantly impacted. The mice that were superinfected at 14-dpii had similar weight loss and survival curves compared to the mice that were singly infected with IAV (Supplementary Table 1 for complete statistical analysis). Thus, susceptibility of mice to secondary *A. fumigatus* infection is maximum during the earlier stages of influenza infection.

**Figure 2.**
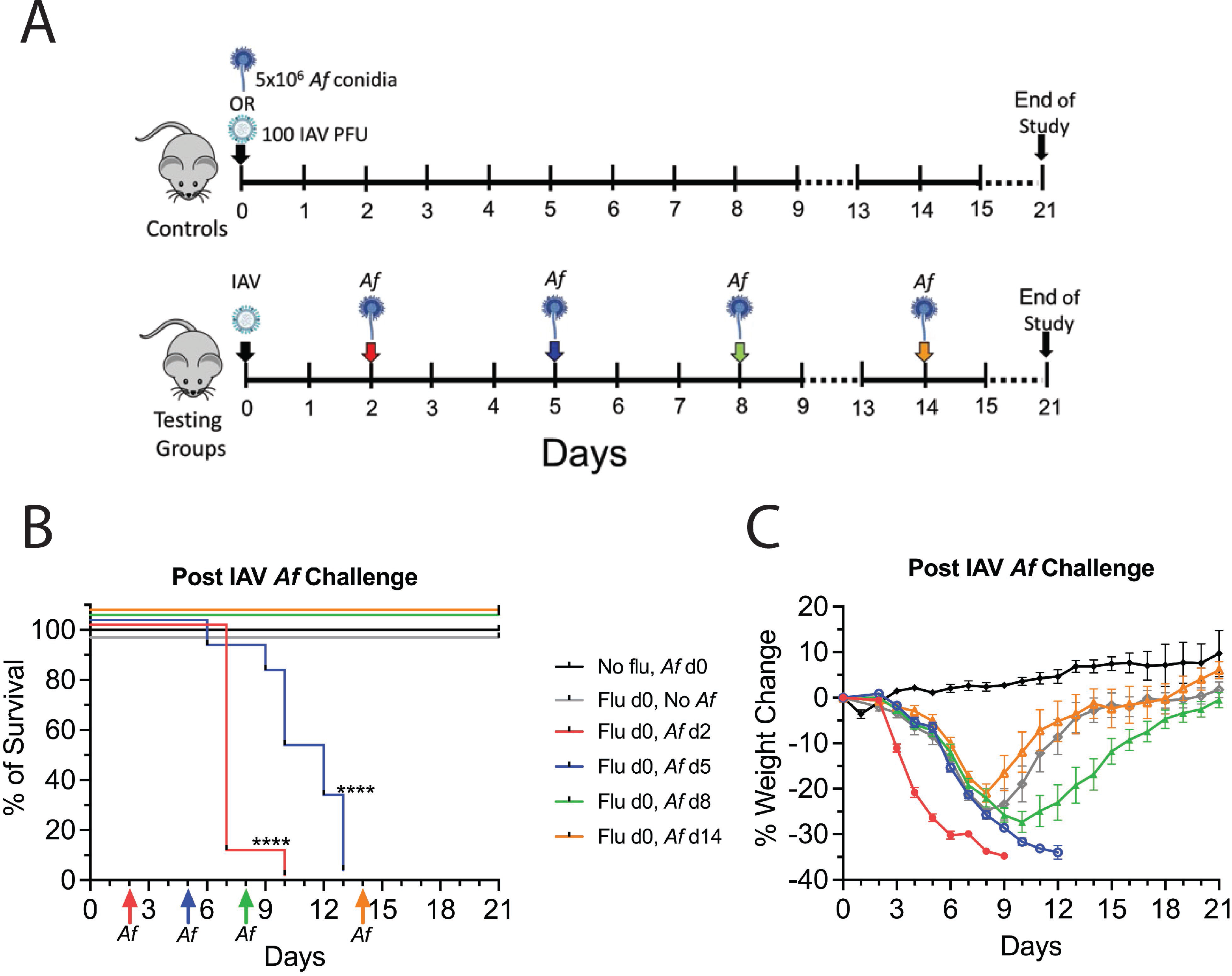
Experimental design for the superinfection model. Mice were first infected with IAV and subsequently challenged with *A. fumigatus*. (A) Schematic description of the experimental design. Mice that were singly challenged with 5×10^6^ *A. fumigatus* CEA10 conidia (O.T.) and singly infected with IAV at 100 PFU/mouse (I.N.) were used as control groups for the experiments. For our superinfection model, mice were infected with IAV on day 0 and were then challenged with *A. fumigatus* CEA10 conidia at 2-, 5-, 8-, and 14-days post IAV infection (dpii). After the mice were infected, their survival (B) and body weight changes (C) were monitored daily for 21 days post-IAV infection. Data were combined from two experiments with N=5 for each experiment except for the single IAV infected control, which was one experiment with N=5. Data in (C) are means ± SEM. **** P<0.0001 compared with mice infected with only IAV or *A. fumigatus* using the Mantel-Cox test. Statistical analysis of the weight change curves is shown in Supplementary Table 1.

### Secondary aspergillosis alters cytokine and chemokine expressions after IAV infection

To examine the hypothesis that excessive or dysregulated inflammatory responses may contribute to the lethality in experimental IAPA, groups of mice were infected with influenza on day 0 and subsequently challenged with *A. fumigatus* at 2-, 5-, 8-, and 14-dpii. Lung samples were then collected at 24 and 48hr after *A. fumigatus* challenge and analyzed for cytokines and chemokines. Control mice were left uninfected, challenged with *A. fumigatus* alone, or infected with IAV alone. For the group infected with IAV alone, lung samples were collected at the same time points as the superinfected mice (Fig. 3A). For each cytokine and chemokine analyzed, a unique patten was discerned. Compared to the control mice, the superinfected mouse lungs altered the cytokine expression profile for IL-6, TNFα, IFNβ, IL-12p70, IL-1α, and IL-1β (Fig 3B). The superinfected lungs had significantly elevated levels of IL-6, TNFα, and IFNβ during early stages of influenza infection, but their levels dropped if the fungal challenge occurred at later stages of influenza infection. The expression levels for IL-12p70, IL-1α, and IL-1β from the superinfected lungs persisted even at later time points. The superinfected mouse lungs also contained higher levels of the chemokines and growth factors CXCL1, G-CSF, MIP-1α, MIP-1β, RANTES, and MCP-1 at their respective time points compared to controls (Fig 4). Lung levels of IL-12p40, IFNɣ, eotaxin, MIP-1α, GM-CSF, IL-2, IL-10, IL-4, IL-13, IL-5, IL-17A, IL-3, and IL-9 were either low or were similar comparing the control and experimental groups (Supplementary Fig. S1). We next looked for evidence of acute systemic sepsis by measuring serum concentrations of the pro-inflammatory cytokines IL-6 and TNFα in mice. Serum levels of IL-6 and TNFα were not significantly different comparing superinfected mice with mice singly infected with IAV or *A. fumigatus* (supplementary Fig. S2A and B).

**Figure 3.**
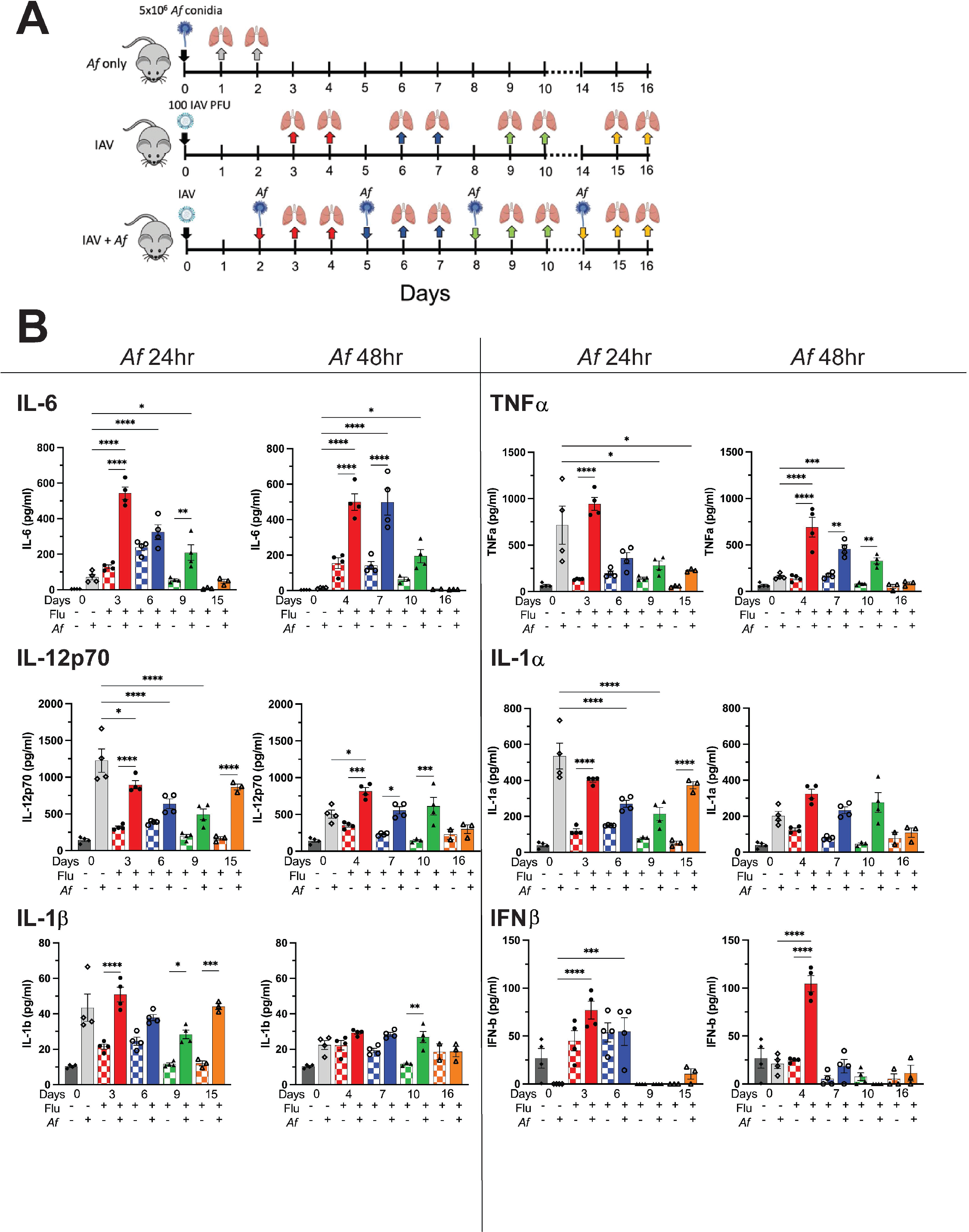
Lung pro-inflammatory cytokine concentrations following IAV and *A. fumigatus* single infections and superinfection. (A) Schematic description of the model. Mice were infected with IAV at 100 PFU/mouse (I.N.) on day 0. The mice were subsequently challenged with 5×10^6^ *A. fumigatus* CEA10 conidia (O.T.) at 2-, 5-, 8-, and 14-dpii. Controls included mice that were uninfected (not shown on the schematic), infected with IAV only, and challenged with *A. fumigatus* only. Lung samples were collected at 24 and 48hr post *A. fumigatus* challenge. Lung samples for control mice singly infected with IAV were collected at the same time points as for the superinfected mice. (B) Cytokine and chemokine levels, as determined by multiplex assay or ELISA on lung homogenates. There were 4 mice per group, except for 14-dpii groups, which had 3 mice per group. Each symbol represents an individual mouse. Data are combined from two independent experiments and expressed as means ± SEM. * P<0.05, ** P<0.005, *** P<0.0005, and **** P<0.0001 by two-way ANOVA with Tukey’s multiple comparison test. Additional cytokines and chemokines are shown in Figure S1 and Figure 4.

**Figure 4.**
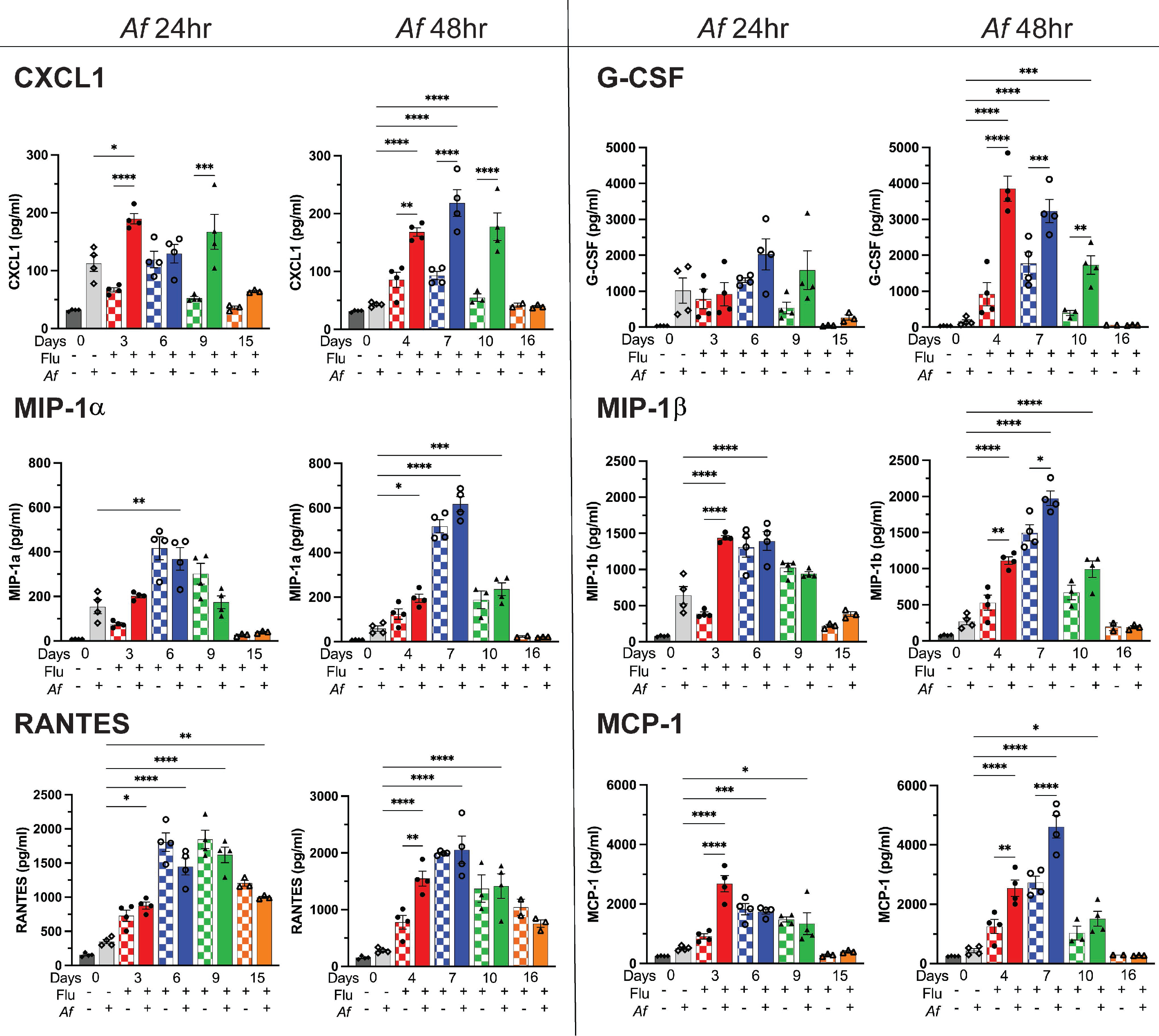
Lung chemokine and growth factor concentrations following IAV and *A. fumigatus* single infections and superinfection. See the Figure 3 legend for details. * P<0.05, ** P<0.005, *** P<0.0005, and **** P<0.0001 by two-way ANOVA with Tukey’s multiple comparison test.

### Histopathological analysis indicates inflammation in IAPA is mostly driven by IAV infection, and secondary aspergillosis does not further increase the inflammation

The results from the multiplex assay showing high levels of pro-inflammatory cytokines in the superinfected mice suggest the lung samples could show extensive inflammation. To investigate this, we performed histopathology of the lungs of mice following IAV and *A. fumigatus* single and dual infections. The protocol was as described in Fig. 3A except lungs were harvested at 24, 72, and 120hr after *A. fumigatus* challenge and sections stained with H&E and GMS. The H&E-stained slides were scored by a blinded pathologist based on the percentage of inflammation in the lung (Fig. 5A). When the mice were singly challenged with *A. fumigatus*, roughly 10% of the lung was inflamed and the inflammation largely dissipated by 120hr after the *A. fumigatus* challenge. In contrast, when the mice were infected with IAV, the percentage of inflammation in the lungs increased as the disease progressed, peaking around 8- to 10-dpii at nearly 50%, and then dropping down to 15% at 19-dpii. The degree of inflammation correlated with the amount of weight loss seen in the previous experiments (Fig. 1B and 2C). Interestingly, the superinfected mice did not show increased inflammation compared to the mice that were singly infected with IAV. However, we observed approximately 30% of the lungs were still inflamed at 120hr after *A. fumigatus* challenge when the IAV-infected mice were challenged with *A. fumigatus* at 14-dpii. Inflammation was localized to lung segments (Supplementary Fig. S3).

**Figure 5.**
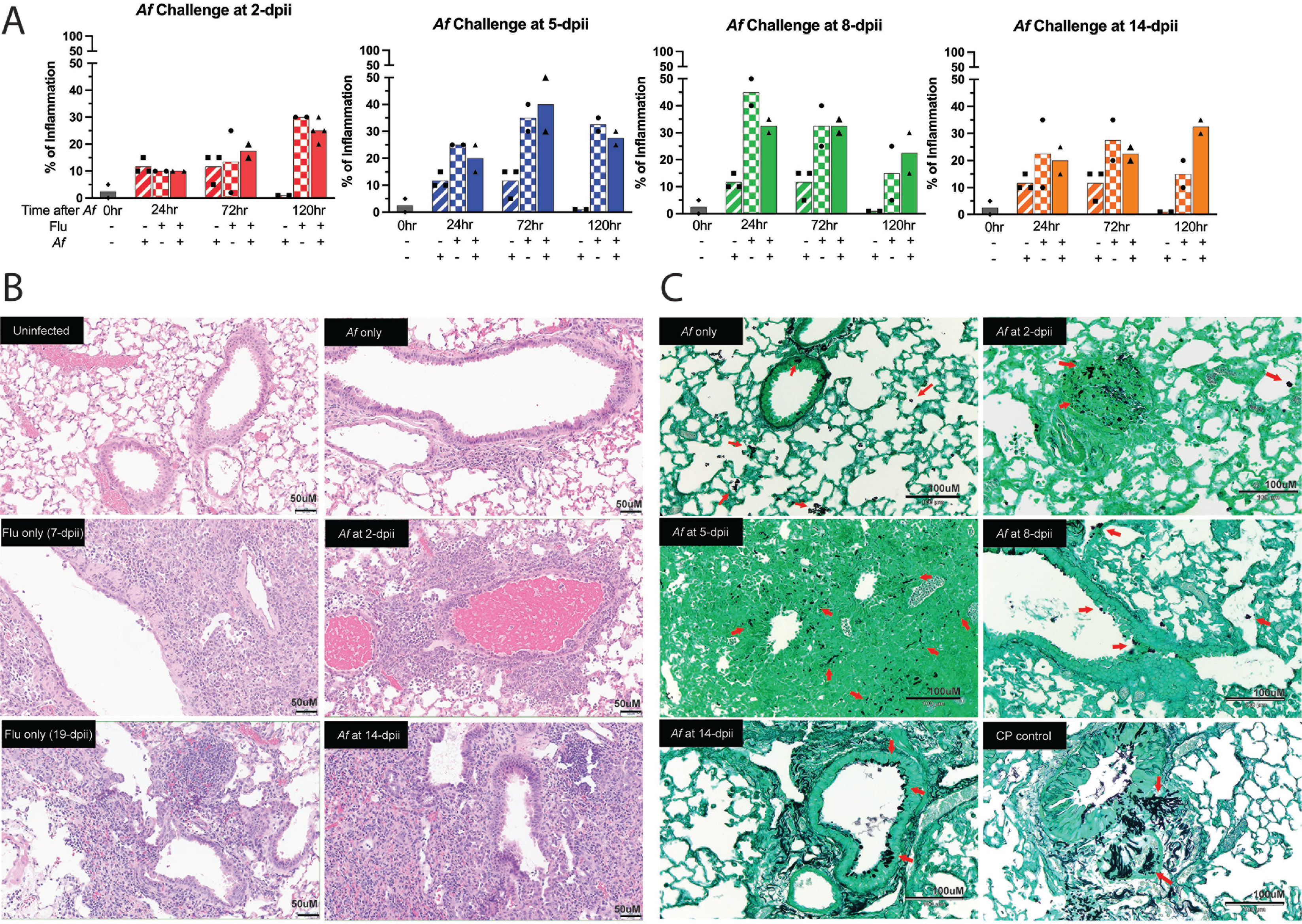
Lung pathology following IAV and *A. fumigatus* single infections and superinfection. Mice were infected with IAV and then challenged with *A. fumigatus* as described in Figure 3A, except lungs were harvested at 24, 72 and 120hr after *A. fumigatus* challenge. Uninfected, *A. fumigatus* challenged only, and IAV infected only were used as controls and collected at the same time points as the superinfected mice. Cyclophosphamide (CP) treated mice infected with *A. fumigatus* served as a positive control for invasive aspergillosis. The data are combined from 5 independent experiments at different time points and each symbol represents an individual mouse. (A) Percentage of inflammation seen on H&E sections of lungs, as determined by a pathologist blinded to the experimental condition. Representative histology of H&E stained at 200X magnification (B) and GMS stained at 20X original magnification (C) of lung samples at 120hr after *A. fumigatus* challenge. Scale bars are 50 microns for (B) and 100 microns for (C). Red arrows point to conidia or hyphae in the GMS-stained lung samples.

Thereafter, we focused on lung samples that were collected at 120hr after *A. fumigatus* challenge as the superinfected mice start to die at this time point. Lungs from mice challenged with just *A. fumigatus* showed sparse chronic inflammatory cells along very few airways. In contrast, lungs that were either singly infected with IAV or dually infected with IAV and *A. fumigatus* had inflammatory cells composed of mostly histiocytes, lymphocytes, and some eosinophils (Fig. 5B). To investigate the fungal burden, 20 GMS-stained lung fields per mouse were scored for the presence of germ tubes or hyphae. Fields containing even a single germinated conidium were scored positive. While conidia in and along the airways and the parenchyma of the lungs were seen, geminated conidia were only observed in lungs from mice that were superinfected at 2- and 5-dpii and studied 120hr post fungal challenge (Fig. 5C and Supplementary Fig. S4). Even in those mice, germination was seen in fewer than 15% of the fields. In comparison, extensive hyphal growth was noted in the lungs of mice treated with cyclophosphamide, a chemotherapy drug that predisposes mice to experimental aspergillosis (20, 21). Taken together, the lung histopathology demonstrates that the superinfected mice do not develop widespread invasive aspergillosis. In addition, *A. fumigatus* challenge after IAV infection does not further increase the percentage of pulmonary inflammation.

### IAV infection does not have a significant impact on fungal clearance

To examine the hypothesis that IAV infection might promote fungal growth during the early stage of IAV infection, we chose a lethal time point (at 2-dpii) and a nonlethal time point (at 14-dpii) to superinfect the IAV-infected mice with *A. fumigatus*. Mice were infected as in Fig. 3A, except lungs were collected at 24, 48, and 120hr after *A. fumigatus* challenge. At the lethal 2-dpii time point, by RT-qPCR, the IAV viral load was high with even higher viral loads at 48hr in the mice that were superinfected with *A. fumigatus* (Fig. 6A), and it remained relatively high even at 120hr after the *A. fumigatus* challenge. We noticed IAV infection had an impact on fungal clearance after *A. fumigatus* challenge especially at 120hr post *A. fumigatus* challenge, as measured by RT-qPCR (Fig. 6B) and CFUs (Fig. 6C). Despite the increased fungal burden in the lungs, extrapulmonary spread of *A. fumigatus* to brain, liver, spleen and kidneys was not observed (supplementary Fig.S5). By 14-dpii, the viral load was undetectable, except for one sample which was just above the lower limit of detection (Fig. 6D). Although there was a delay in fungal clearance at 48hr after *A. fumigatus* challenge as assessed by conidial equivalents (Fig. 6E) and CFUs (Fig. 6F), the fungal burden returned to the baseline level by 120hr post *A. fumigatus* challenge.

**Figure 6.**
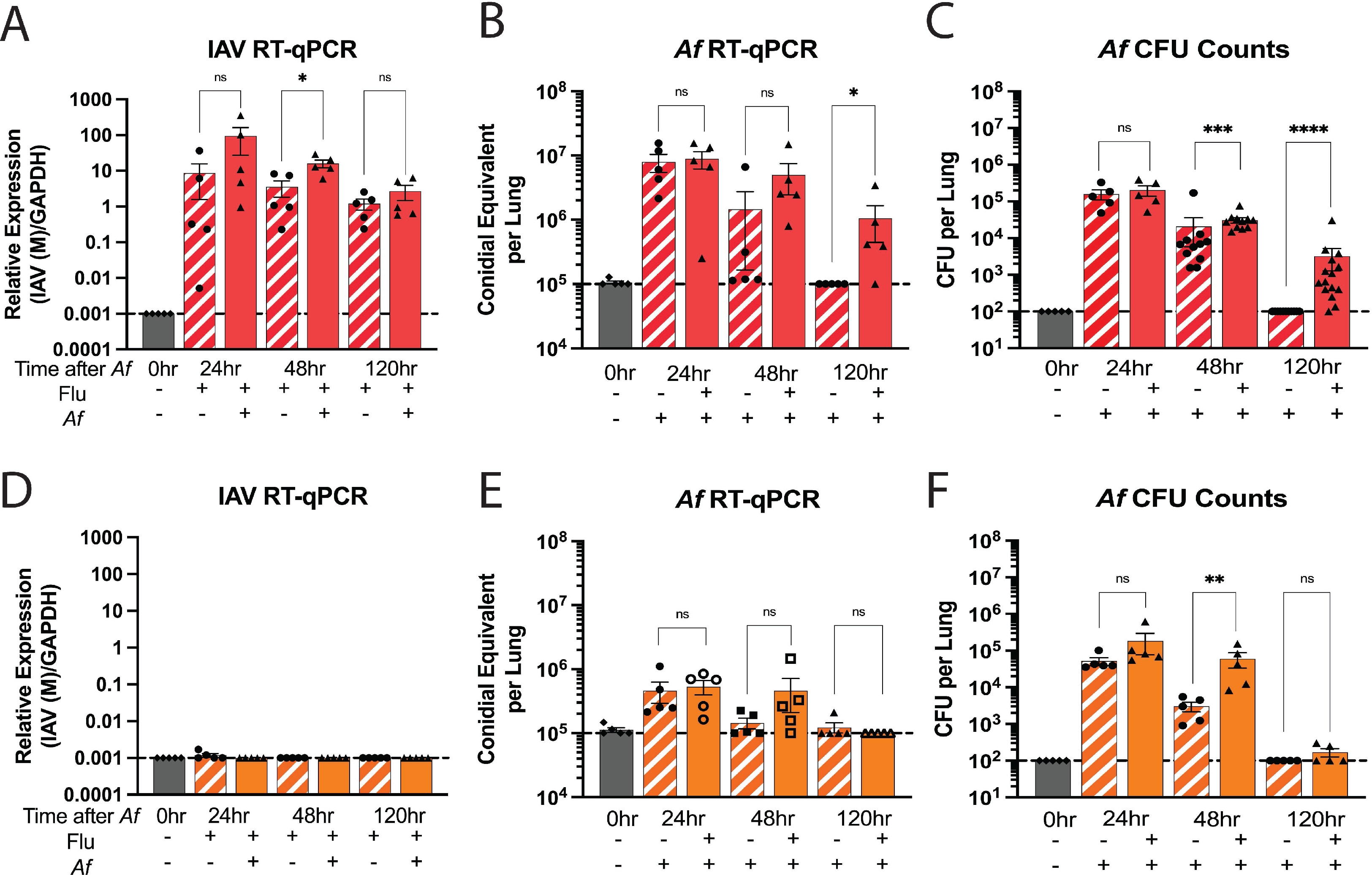
Viral and fungal load in the lungs as determined by RT-qPCR analysis and CFU plating. Mice were infected I.N. with 100 PFU IAV/mouse. At either 2-dpii (red) or 14-dpii (orange), mice were then challenged O.T. with 5×10^6^ *A. fumigatus* CEA10 conidia. Lung samples were collected at 24, 48, and 120hr after *A. fumigatus* challenge. Uninfected, single IAV infected, and single *A. fumigatus* challenged mice were used as controls. Lung samples from the control mice were collected at the respective time point of the superinfected mice. There are 5 mice per group. The data shown are the combination of 4 independent experiments at different time points and expressed as means ± SEM and each symbol represents an individual mouse. Top portion: Mice were challenged with *A. fumigatus* at 2-dpii; the relative expression of influenza (A), *A. fumigatus* conidial equivalent (B), and *A. fumigatus* CFU counts (C) were measured by RT-qPCR or CFU plating. Bottom portion: Mice were challenged with *A. fumigatus* at 14-dpii and the relative expression of influenza (D), *A. fumigatus* conidia equivalent (E), and *A. fumigatus* CFU counts (F) were also measured by RT-qPCR or CFU plating. Dotted line represents the lower limit of detection (LLD) of the assay. Individual data points, means, and SEM are shown; numbers at or below the LLD were assigned the value of the LLD. Statistics were calculated using the Mann-Whitney (nonparametric) t test with Bonferroni’s correction for multiple comparisons. * P<0.05, ** P<0.005, *** P<0.0005, and **** P<0.0001.

### Neutrophil, but not macrophage, recruitment is dampened when mice are challenged with *A. fumigatus* during the early stage of IAV infection

In the final sets of experiments, we used fluorescent *Aspergillus* reporter (FLARE) conidia (18, 22, 23) combined with flow cytometry analysis to examine leukocyte recruitment to the lungs as well as fungal uptake and killing by phagocytic cells. FLARE conidia are genetically encoded with dsRed and then stained with the secondary fluorophore Alexa Fluor 633 (AF633). Conidia killed by phagocytes lose their dsRed staining but retain their AF633 staining (Supplementary Fig S6 and gating strategy: Supplementary Fig. S7). This enables FLARE conidia to be used as a tool to distinguish live conidia (dsRed^+^ and AF633^+^) from dead conidia (dsRed^-^ and AF633^+^). Moreover, when combined with labeled antibodies that distinguish neutrophils and macrophages, phagocytosis and killing of FLARE conidia by these cell types can be assayed. Three caveats need to be considered when interpreting flow cytometric assays with FLARE conidia. First, FLARE conidia that germinate lose dsRed expression. Second, phagocytic cells that contain multiple FLARE conidia cannot be distinguished from cells with just one conidium. Finally, cells with a mixed population of phagocytosed live and dead FLARE conidia will be positive for dsRed and AF633 and thus be scored as having just live conidia.

For the first set of experiments with FLARE conidia, mice were infected with IAV on day 0 and challenged with 3×10^7^ FLARE conidia on 2-, 5-, 8-, and 14-dpii. A higher dose of FLARE conidia was used compared with previous experiments to increase the sensitivity of the flow cytometry assay to detect phagocytosed conidia. Control groups consisted of mice that were uninfected, infected with IAV alone, or challenged with *A. fumigatus* FLARE conidia alone. After the mice were challenged with FLARE conidia, lung samples were collected 24hr post challenge. Lung samples from the single IAV infected mice were collected at the respective time points of 3-, 6-, 9-, and 15-dpii (Fig. 7A). In terms of total leukocyte counts in the lung, as determined by expression of CD45^+^, no differences between superinfected mice and single IAV infected mice were observed at any time point. However, leukocyte numbers were higher in superinfected mice compared to mice challenged with *A. fumigatus* alone at 8- and 14-dpii (Fig. 7B). Neutrophil recruitment was mainly driven by the *A. fumigatus* challenge as the mice that were singly infected with IAV had significantly fewer neutrophils recruited to the lung. Compared with mice that were challenged with FLARE conidia alone, neutrophil recruitment to the lungs of dually infected mice was impaired during the early, but not late, stages of IAV infection. Interestingly, the total number of neutrophils was higher when the mice were challenged with FLARE conidia at 8-dpii (Fig. 7C). As for the macrophage population, significant lung recruitment was not observed in any of the groups (Fig. 7D).

**Figure 7.**
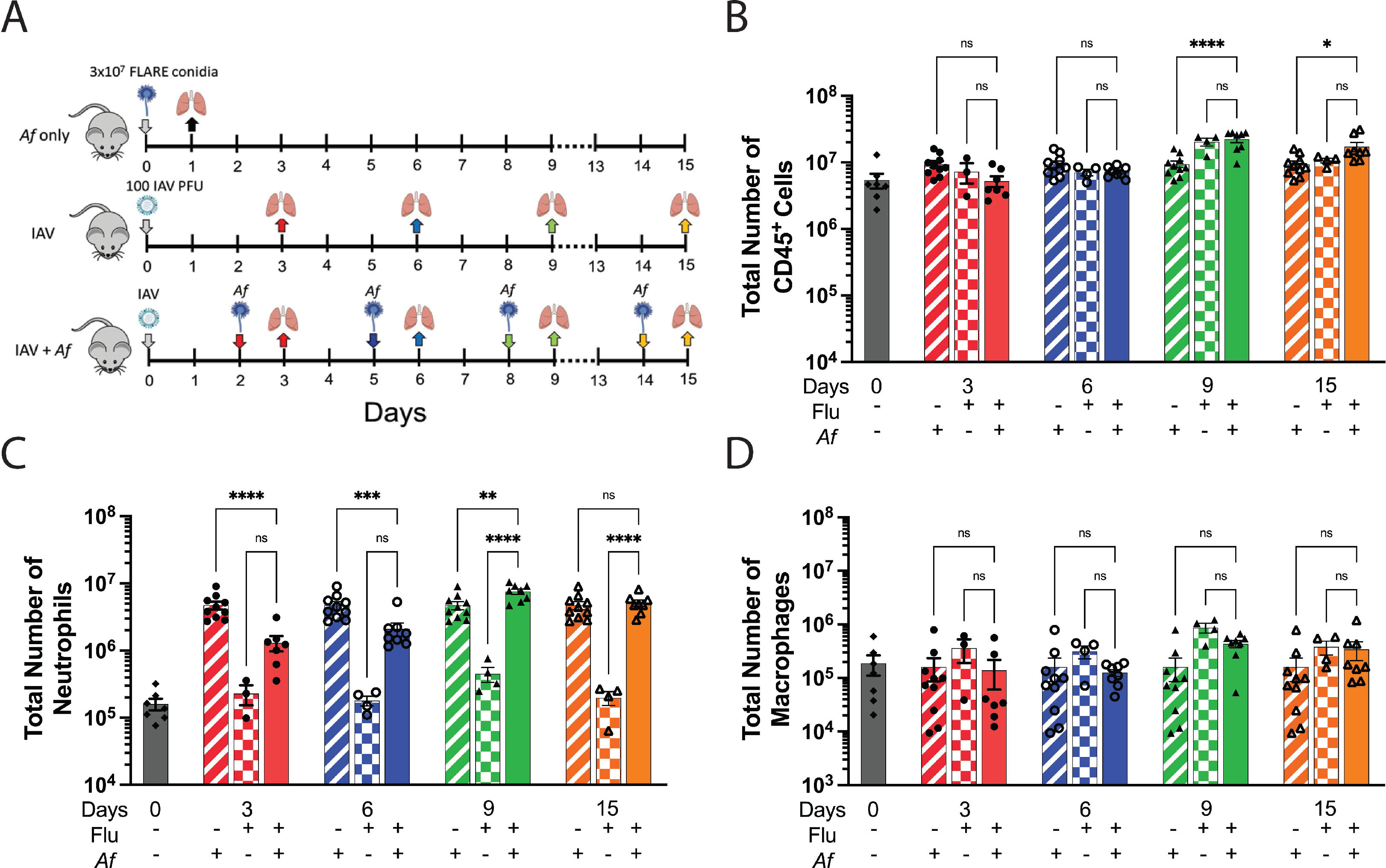
Flow cytometric analysis of leukocyte populations in lung samples from mice following IAV and *A. fumigatus* single infections and superinfection. (A) Schematic description of the experiment. Uninfected mice, singly *A. fumigatus* (*Af*) FLARE conidia challenged mice, and singly IAV infected mice were used as controls. Mice were infected I.N. with 100 PFU IAV followed by 3×10^7^ FLARE conidia O.T. challenge at 2-, 5-, 8-, and 14-dpii. Lung samples were collected at 24hr post FLARE conidia challenge. For the control mice, lung samples were collected at the respective time points of the superinfected mice. Supplementary Figure S4 shows the gating strategy. (B, C, and D) Number of leukocytes (CD45^+^), neutrophils (CD45^+^, CD11b^+^, Ly6C^+^, Ly6G^+^, F4/80^-^), and macrophages (CD45^+^, CD11b^+^, Ly6G^-^, Ly6C^+/-^, F4/80^+^), respectively, in the lungs as a function of type of infection and time after infection. The data shown are the combination of ≥5 independent experiments at different time points and each symbol represents an individual mouse. * P<0.05, ** P<0.005, *** P<0.0005, and **** P<0.0001 by two-way ANOVA with Tukey’s multiple comparison test. ns, not significant.

### Influenza infection does not have a major effect on phagocytosis and fungal killing of *A. fumigatus* conidia

We then examined *A. fumigatus* phagocytosis and killing by lung neutrophils and macrophages following challenge of mice with FLARE conidia. The neutrophil and macrophage populations shown in Fig. 7 were further analyzed for the presence of live or dead FLARE conidia using the gating strategy shown in supplementary Fig. S7. Compared with mice not infected with IAV, the percentage of lung neutrophils containing conidia was significantly higher during the early stages of IAV infection (Fig. 8A), suggesting IAV infection did not suppress the ability of neutrophils to phagocytose conidia. Interestingly though, the total number of neutrophils containing conidia was similar during different stages of IAV infection (Fig. 8B). Thus, mice could compensate for having fewer neutrophils recruited to the lung (Fig. 7C) by increasing the percentage of neutrophils phagocytosing conidia. We observed the same trends for fungal killing by neutrophils. There was a higher percentage of neutrophils containing dead conidia (dsRed^-^ and AF633^+^) at the early stages of influenza infection (Fig. 8C), but there were no differences observed between the groups in terms of the total number of neutrophils containing dead conidia (Fig. 8D).

**Figure 8.**
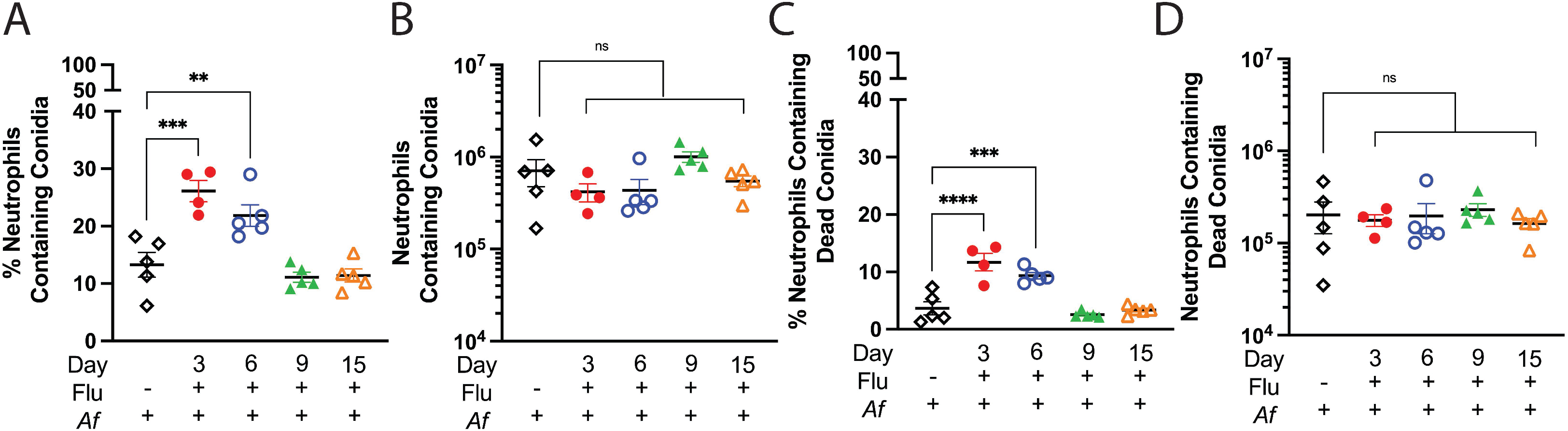
Phagocytosis and killing of *A. fumigatus* FLARE conidia by neutrophils. Mice were infected with FLARE conidia at 2, 5, 8, and 14 dpii, as described in Figure 7A. Lungs were harvested at 24hr, single cell suspensions were made, and the neutrophils analyzed by flow cytometry for the presence of live and dead FLARE conidia according to the schematic in Figure S4. A control group of mice was challenged with FLARE conidia alone (no IAV). (A) and (B) The percentage of neutrophils containing conidia and the total number of neutrophils with conidia, respectively. (C) and (D) Percentage of neutrophils and total number of neutrophils containing only dead conidia, respectively. ** P<0.005, *** P<0.0005, and **** P<0.0001 by one way ANOVA comparing the superinfected groups with the mice singly challenged with *A. fumigatus*, ns, not significant. Data are the combination of ≥5 independent experiments at different time points and each symbol represents an individual mouse.

Compared to neutrophils, macrophages contributed less to fungal uptake and killing. On average, less than 10% of macrophages had phagocytosed conidia (Supplementary Fig. S8A), and the total number of macrophages containing conidia was approximately ten times lower than those of the neutrophils (Supplementary Fig. S8B). Fungal killing by macrophages was not significantly affected by IAV infection, regardless of the time point studied (Supplementary Fig. S8C-D). A portion from each of the lung homogenates was plated on Sabouraud dextrose agar; no differences in fungal burden, as measured by CFUs, were observed comparing control and experimental groups (Supplementary Fig. S9).

## Discussion

Patients with severe influenza are at risk for secondary bacterial and fungal infections, including invasive pulmonary aspergillosis (12–14). In a multicenter cohort study, invasive aspergillosis was diagnosed in 19% of patients with an intensive care unit admission diagnosis of influenza (14). Herein, we developed a mouse model to examine mechanisms by which influenza infection predisposes to superinfection with *A. fumigatus*. In contrast to other published mouse models (15–17, 19) which used inocula of *A. fumigatus* ranging from 2.5 – 10 × 10^7^ conidia, we used 5.0 × 10^6^ conidia reasoning that the lower inoculum would be less likely to overwhelm the mouse and perhaps be more physiological. Nevertheless, in agreement with investigators who used higher inocula, we found that mice infected with a sublethal dose of IAV had 100% mortality following a subsequent challenge with *A. fumigatus*. However, the timing of the superinfection was critical as 100% mortality was seen in mice superinfected during the early stages (days 2 and 5) of IAV infection whereas all mice challenged at later time points (days 8 and 14) survived (Fig. 2).

In contradistinction to studies which used higher fungal inocula (15, 16, 19), widespread hyphal growth was not observed in our experiments (Fig. 5C and Supplementary Fig, S4). Most lung specimens contained no germinated conidia or hyphae on histopathology. The only lung samples that exhibited germinated conidia were the superinfected mice that had prior IAV infection at 2 or 5 days and were examined at 120hr post *A. fumigatus* challenge. Based on the mortality curves (Fig. 2C), this is shortly before the mice die. The histopathology findings are supported by RT-qPCR and CFU analyses which found modestly increased fungal burdens in the dually infected mice at the time point before predicted death (Fig. 6). Taken together, our data suggest that while fungal burden could have contributed to the mortality seen during dual infection with IAV and *A. fumigatus*, it is unlikely to be the primary cause of death as extensive hyphal growth in the lungs or extrapulmonary spread of *A. fumigatus* (supplementary Fig. S5) was not seen in the superinfected mice.

Neutrophils play a crucial role in host defenses against *A. fumigatus* (1, 2). In a different model of *A. fumigatus* superinfection following IAV infection, mice infected with influenza infection had decreased pulmonary concentrations of the neutrophil chemoattractants CXCL1 and CXCL2 following *A. fumigatus* challenge (15). Although the superinfected mice were not neutropenic, they did have reduced neutrophil recruitment to the lungs compared with mice infected with just *A. fumigatus*. We also identify that IAV infection had a suppressive effect on neutrophil recruitment following fungal challenge. However, reduced numbers of lung neutrophils were only seen in mice superinfected with *A. fumigatus* during early stages (day 3 and 6) of the IAV infection, but not during the later stages (day 9 and 15) (Fig. 7C). In addition, at most of the time points studied, superinfected mice had significantly higher lung CXCL1 levels compared with mice infected with *A. fumigatus* alone (Fig. 4).

Liu *et al*. observed neutrophil recruitment to the lungs 36hr post *A. fumigatus* challenge was not affected by influenza infection 6 days prior to fungal challenge (16). They did, however, find inhibitory effects of IAV infection on conidial phagocytosis, phagolysosome maturation, and conidial killing. Liu *et al*. did not find diminished production of reactive oxygen species by neutrophils and macrophages, which others have found when IAV-infected phagocytes were superinfected with *Staphylococcus aureus* and *Cryptococcus gattii* (24, 25). We did not observe significant impairments in neutrophil phagocytosis and conidial killing at 24hr after *A. fumigatus* challenge in IAV-infected mice (Fig. 8). Moreover, at the four timepoints studied, IAV had no effect on F4/80^+^ pulmonary macrophages in terms of cell numbers (Fig 8), and conidial phagocytosis and killing (Fig. S6). Thus, in our model, impairment of neutrophil and macrophage recruitment and phagocyte fungicidal activity does not appear to explain the lethality in mice with IAPA. We speculate the disparate results are likely a result of differences in experimental models and emphasize the importance of studying multiple time points to get a full picture of the immunology of IAPA. The translational importance of studying early timepoints is evidenced by a multicenter clinical study of critically ill influenza patients in which 15 out of 21 patients (71%) who developed IAPA did so within 48hr of intensive care unit admission (26).

Recently, Sarden *et al*. showed that natural IgG antibodies produced by innate B1a cells contribute to host resistance to invasive aspergillosis by promoting neutrophil opsonophagocytosis (17). IAV infection of mice led to a depletion of B1a cells and, in an IAV/*A. fumigatus* superinfection model, serum from WT, but not B cell-deficient mice improved survival following pulmonary challenge with conidia. While we did not directly examine the effects of IAV infection on B cells and humoral defenses, we found no significant effects of IAV infection on opsonophagocytosis of *A. fumigatus* conidia. *A. fumigatus* conidia and hyphae activate the alternative complement pathway and are recognized by innate pattern recognition receptors (27–29), which we speculate could compensate for the lack of opsonic antibody.

Cytokines responses can be beneficial to host defenses, but in excessive can be detrimental such as during a “cytokine storm” when a life-threatening, dysregulated cytokine response occurs (30, 31). We examined concentrations of 24 cytokines and chemokines at eight different time points in mice singly or dually infected with IAV and *A. fumigatus* (Figs. 3, 4, and Supplementary Fig. S1). During infection, it can be difficult to distinguish if an elevated cytokine response is beneficial or harmful to the host. Thus, we were particularly interested in cytokines and chemokines that were highly expressed in mice superinfected at time points in which they succumbed to the dual infection. The most striking finding was with IL-6 which was highly expressed in the superinfected mice, particularly in mice superinfected with *A. fumigatus* at the early, lethal time points (Fig. 3B). Future studies are needed to see if blocking IL-6 can reverse the mortality of superinfected mice. If so, then trials of adjuvant IL-6 blockade in selected patients with IAPA might be considered. This type of immunomodulatory approach has clinical precedent; in a trial of hospitalized COVID-19 patients with hypoxia and systemic inflammation, subjects who received the IL-6 blocker tocilizumab had improved outcomes (32).

Although the effects were not as consistent as with IL-6, lung concentrations of the pro-inflammatory cytokines TNFα, IFNβ, IL-12p70, IL-1α, and IL-1β, and the chemokines CXCL1, G-CSF, MIP-1α, MIP-1β, and MCP-1 were also significantly elevated in superinfected mice at one or more early time points compared with mice singly infected with *A. fumigatus* (Figs. 3B, 4). However, elevated levels of IL-6 and TNFα were not found in the serum of the superinfected mice, suggesting systemic sepsis is not a major contributor to death of the superinfected mice (supplementary Fig. S2). Experiments examining variables such as pulmonary function and body temperature changes are necessary though to determine the relative roles of the local and systemic responses to mortality. An interesting observation from the cytokine data are the spikes in lung levels of IL-10, IFNɣ, and eotaxin around 6- to 7-dpii (Supplementary Fig. S1). These spikes were not augmented by superinfection with *A. fumigatus* but do align with one of the lethal time points in our superinfection model (*A. fumigatus* challenge at 5-dpii). van der Sluijs *et al*. reported mice succumbed to secondary *S. pneumoniae* challenge at 14-dpii with elevated levels of IL-10; treatment with neutralizing antibodies against IL-10 reversed the lethal outcome (33). However, in our study, mice challenged with *A. fumigatus* at 14-dpii had 100% survival, suggesting that IL-10 may have a different role in the setting of IAPA. The expression level of other cytokines associated with anti-inflammatory responses and tissue repair, including IL-2, IL-4, IL-5, and IL-13 were low and not significantly altered after IAV infection or *A. fumigatus* challenge. Taken together, the cytokine and chemokine data suggest that post-IAV aspergillosis is associated with mostly pro-inflammatory cytokine responses, especially at the early, lethal time points. While we speculate this pro-inflammatory response is detrimental to the host as it is associated with mortality, cytokine neutralization studies are needed to prove causality.

IAV infection can have severe physical and immunological impact on the host respiratory track including disruption of the mucosal layer, epithelial cell and alveolar damage, edema, tissue necrosis, and cytokine and chemokine dysregulation (9–11). This damage creates a microenvironment that increases susceptibility to secondary infections. However, by histopathological examination, RT-qPCR analysis, and CFU plating, mice challenged with *A. fumigatus* alone almost completely cleared the fungus and had minimal inflammation at the 120hr time point (Fig. 5 and 6). In contrast, IAV infection induced considerably more profound and longer lasting inflammation in the lung tissues. Moreover, despite the IAV viral load in the lungs becoming undetectable by 15-dpii (Fig. 6D), approximately 10% of the lungs remained inflamed by 19-dpii (Fig. 5A). Unexpectedly, secondary aspergillosis did not significantly increase the percentage lung inflammation compared with that seen with IAV infection alone (Fig. 5A). Based on these observations, it appears the inflammation in IAPA is mostly driven by the IAV infection.

Neuraminidase inhibitors, particularly oseltamivir, are routinely used in patients with severe influenza. Interestingly, Dewi *et al.* (34) showed decreased survival of mice treated with oseltamivir and then infected with *A. fumigatus*. The mechanistic basis for the reduced survival was postulated to be inhibition of host neuraminidases leading to diminished immune responses to *A. fumigatus*. These data suggested that treatment of influenza with oseltamivir may predispose patients to secondary aspergillosis. However, when the effects of oseltamivir was studied in a mouse model of IAPA, early treatment of oseltamivir was protective, as assessed by reduced lung damage, diminished severity of influenza infection, and increased survival (19). These results support the notion that the inflammatory response induced by influenza directly predisposes to secondary aspergillosis.

In conclusion, we found vulnerability to secondary aspergillosis is maximal during early stages of influenza infection. While the cause of the high mortality rate in our mouse model of IAPA remains speculative, the data suggest it is likely multifactorial. First, mice infected with influenza alone lost about a quarter of their body weight during the first 9 days of infection. This presumably left the mice dehydrated, nutritionally depleted, and with a reduced capacity to withstand a subsequent *A. fumigatus* challenge, even at a relatively low inoculum that in and of itself leads to minimal weight loss. Second, although the percent lung inflammation was similar when comparing mice infected with IAV alone to mice superinfected with IAV and *A. fumigatus*, the nature of the inflammatory response was different. Many pro-inflammatory cytokines and chemokines were significantly higher in the superinfected mice. Moreover, differences between groups were noted when examining pathology. A surprising finding though was fungal growth did not appear to be a major contributor to death in IAPA. If our findings are confirmed in humans, they suggest a rationale for clinical studies examining the benefit of adjuvant anti-inflammatory agents in the treatment of IAPA.

## Materials and Methods

### Mice

Six weeks old WT C57BL/6 mice of both sexes were purchased from Taconic Biosciences (Rensselaer, NY) and housed in the animal facility at University of Massachusetts Chan Medical School (UMCMS). Mouse experiments were conducted under a protocol approved by the Institutional Animal Care and Use Committee at UMCMS, in accordance with the guidelines from the National Institutes of Health’s Guide for the Care and Use of Laboratory Animals. Infected mice were monitored for body weight and survival daily. Moribund mice were humanely euthanized.

### Influenza A Virus Stock and Mouse Infection

Influenza A/PR/8/34 (PR8), grown in chicken egg allantoic fluid, was purchased from Charles River Laboratory (Wilmington, MA). The stock was titered by viral plaque assay using Madin-Darby canine kidney cells (35). The inocula were made by diluting the influenza stock with sterile PBS (Gibco). Mice were anesthetized with isoflurane and then infected intranasally (I.N.) with doses of PR8 ranging from 5 to 2500 PFU in 50 µL of sterile PBS.

### *Aspergillus fumigatus* Strains, Culture and Mouse Challenge

*A. fumigatus* CEA10 and FLARE conidia were prepared as described (18, 22, 36). Conidia were suspended at the desired concentration in PBS with 0.05% Tween 20. Isoflurane-anesthetized mice were challenged with conidia (50 µL of the fungal suspension) via the orotracheal (O.T.) route. Lungs were harvested at specified time points after influenza infection and/or *A. fumigatus* challenge depending on the experiments. Control mice were left uninfected or challenged with only *A. fumigatus* or IAV.

### Multiplex and ELISA Analysis

Lung samples were collected at specified time points and stored at -80°C. The lung samples were processed by homogenizing with 1x cOmplete Mini protease inhibitor cocktail (Roche) containing 0.05% Triton X-100 (EMD Millpore) and incubated on ice for an hour. Supernatants were collected following centrifugation of the homogenate at 18,000 gravity for 20 minutes at 4°C and then stored at -80°C until analysis by Bio-Plex Pro Mouse Cytokine 23-plex Assay (Bio-Rad Laboratories) following the manufacturer’s protocol. TNFα, IFNβ, and IL-6 (serum) were analyzed using Mouse DuoSet ELISA kits (R&D Systems).

### Histopathology Staining

Lung samples were collected at specific times. As a positive control for invasive aspergillosis, a group of mice received 10 µL/g of intraperitoneal cyclophosphamide (20 mg/mL) (EMD Millipore) every 48hr starting one day before *Aspergillus* challenge. Following euthanasia, lungs were inflated with 500 µL of 3.7% buffered formalin (Fisher brand, Fisher HealthCare) through the trachea and then removed immediately from the mouse. The collected lungs were put into a pathology cassette and stored in buffered formalin at room temperature until processing. The lung samples were sent to the Morphology Core Facility at UMCMS (https://www.umassmed.edu/morphology/) where 5 µm thick sections were cut and stained with hematoxylin and eosin (H&E) or Grocott’s methenamine silver (GMS) following the manufacturer’s protocol (Stat Lab). The H&E-stained sections were then read in a blinded fashion by a pathologist. Slides were scored for the percentage of inflammation and analyzed with regards to the type of infiltration. Fungal morphology and fungal burden were determined by reading the GMS-stained slides. The percentage of microscope fields with germinated conidia was determined by randomly selecting 20 lung fields from the GMS-stained samples at 20X magnification. Fields that contained at least one germinated conidia were scored as positive.

### CFU and RT-qPCR analyses

Lungs were harvested and placed in one mL of PBS containing 0.05% Tween 20 and 1 mg/mL DNase I (Sigma). The tissue was homogenized using C-Tubes from Miltenyi Biotech. To determine CFUs, 10 µL were removed, diluted in UltraPure distilled water (Invitrogen) and plated on Sabouraud dextrose agar (Remel). The plates were incubated at room temperature for 3 to 5 days at which time the colonies became visible to count.

To perform RT-qPCR analysis, zirconia beads (0.6g, BioSpec Products) were added to the lung samples and homogenized with a Mini-Beadbeater-8 (BioSpec) for 2.5 minutes at room temperature. After homogenization, samples were stored at -80°C until RNA extraction. A 200 µL portion of the sample was taken for RNA extraction using TRIzol Reagent following the manufacturer’s protocol. KAPA SYBR FAST One-Step universal kit (Roche) and LightCycler 96 (Roche) were used to conduct and analyze the RT-qPCR for all the RNA samples. Primers used in the experiments are listed on Table 1. Conidial equivalent was calculated by comparing the cycle threshold (Ct) value of each sample to a standard curve generated by adding known amounts of CEA10 conidia (1×10^3^ to 1×10^8^) to WT mouse lungs. The Ct values from the influenza and glyceraldehyde-3-phosphate dehydrogenase (GAPDH) samples were converted into gene number with the equation: 10^((Ct minus number of cycles)/-3.32). Then the gene number of the IAV matrix protein was divided by the gene number of GAPDH to determine relative expression.

**Table 1.**
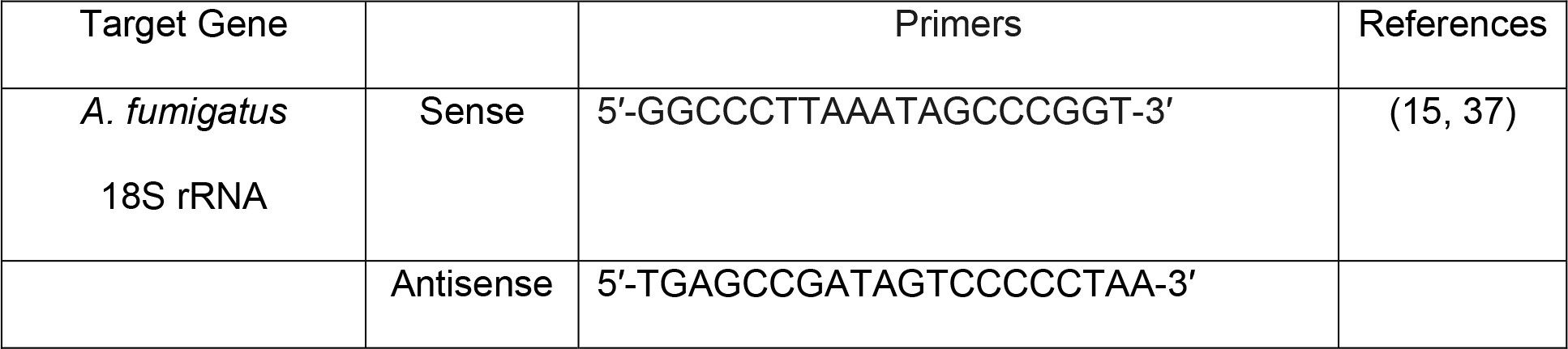
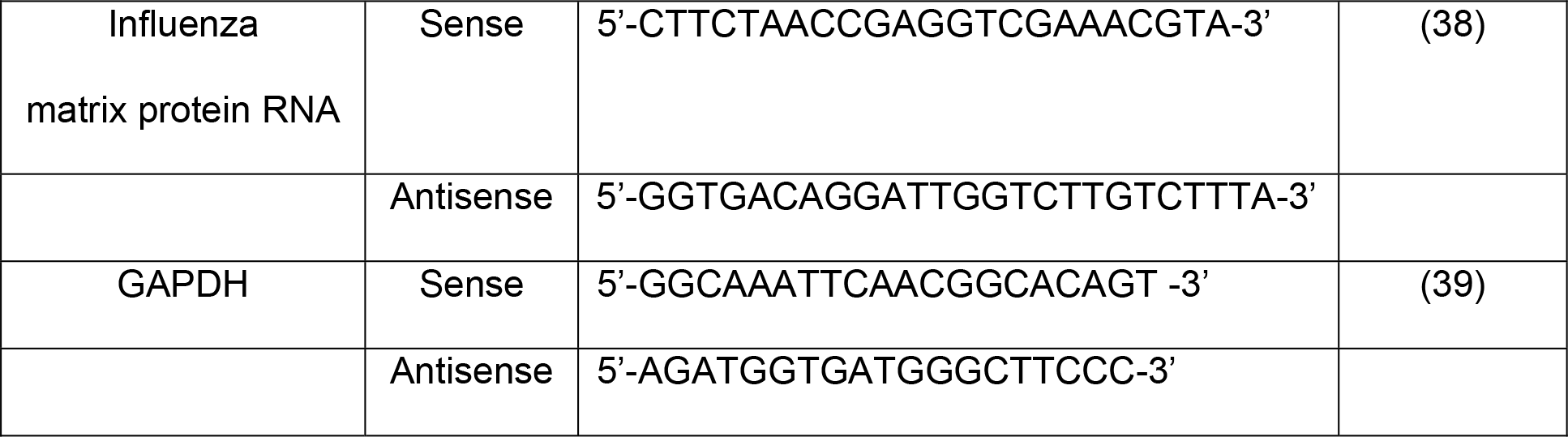
List of Primers for RT-qPCR assays.

### Flow Cytometry Analyses

Lungs were harvested, cut into small pieces with scissors, and resuspended with 5 mg/mL collagenase and 1 mg/mL DNase I in 1 mL of RPMI 1640 without phenol red (Gibco). After incubation at 37°C for a half hour, the lung samples were meshed with the plunger of a syringe on a 70µm cell strainer and rinsed with RPMI 1640 without phenol red. To determine CFUs, 10 µL were removed and plated as described above.

To analyze the lung cells by flow cytometry, red blood cells (RBC) were lysed with RBC buffer (Invitrogen), and the remaining cells were washed and resuspended in 1 mL of RMPI 1640 without phenol red. Then, 100 µL of the sample were stained for cell surface antigens. First, Fc receptors were blocked with anti-mouse CD16/CD32 monoclonal antibody 2.4G2 (BD Pharmingen) following the manufacturer’s instruction. Surface markers were stained with antibodies listed in Table 2. Flow cytometer data were acquired with a 5-Laser Cytek Aurora cytometer and analyzed with FlowJo X software (Tree Star Inc.)

**Table 2.** List of surface marker antibodies.

| Monoclonal Antibodies | Clone | Dilution | Manufacturer |
| --- | --- | --- | --- |
| CD45 – BV785 (0.2 mg/mL) | 30-F11 | 1:200 | Biolegend |
| CD11b – AF700 (0.2 mg/mL) | M1/70 | 1:200 | Invitrogen |
| CD11c – BV650 (0.2 mg/mL) | N418 | 1:200 | Biolegend |
| Ly-6C – PE-Cy7 (0.2 mg/mL) | HK1.4 | 1:100 | Biolegend |
| Ly-6G – PerCP-Cy5.5 (0.2 mg/mL) | 1A8 | 1:100 | Biolegend |
| F4/80 – APC-Cy7 (0.2 mg/mL) | BM8 | 1.5:100 | Biolegend |

### Graphs and Statistics

Graphs were generated and statistics calculated using Prism GraphPad Software (version 9.4). Graphic designs were created using BioRender.

### Data Availability

Materials and data that are reasonably requested will be made available in a timely fashion to members of the scientific community for noncommercial purposes.

## ACKNOWLEDGMENTS

This research was supported by the National Institute of Allergy and Infectious Diseases, National Institutes of Health grants R01 AI139615, R01 AI172154, and R01 AI125045. The authors thank Robert Cramer (Geisel School of Medicine, Hanover, NH) and Tobias Hohl (Weill Cornell Medical, New York, NY) for the gifts of *A. fumigatus* CEA10 and FLARE strains, respectively.

## References

1. Latge JP, Chamilos G. 2019. Aspergillus fumigatus and Aspergillosis in 2019. Clin Microbiol Rev 33:e00140–18.

2. Dagenais TR, Keller NP. 2009. Pathogenesis of Aspergillus fumigatus in Invasive Aspergillosis. Clin Microbiol Rev 22:447–65.

3. Park SJ, Mehrad B. 2009. Innate immunity to Aspergillus species. Clin Microbiol Rev 22:535–51.

4. Bongomin F, Gago S, Oladele RO, Denning DW. 2017. Global and Multi-National Prevalence of Fungal Diseases—Estimate Precision. Journal of Fungi 3:57.

5. Brown GD, Denning DW, Gow NA, Levitz SM, Netea MG, White TC. 2012. Hidden killers: human fungal infections. Sci Transl Med 4:165rv13.

6. Koehler P, Bassetti M, Kochanek M, Shimabukuro-Vornhagen A, Cornely OA. 2019. Intensive care management of influenza-associated pulmonary aspergillosis. Clin Microbiol Infect 25:1501–1509.

7. Vanderbeke L, Spriet I, Breynaert C, Rijnders BJA, Verweij PE, Wauters J. 2018. Invasive pulmonary aspergillosis complicating severe influenza: epidemiology, diagnosis and treatment. Curr Opin Infect Dis 31:471–480.

8. Heltzer ML, Coffin SE, Maurer K, Bagashev A, Zhang Z, Orange JS, Sullivan KE. 2009. Immune dysregulation in severe influenza. J Leukoc Biol 85:1036–43.

9. Vogel AJ, Harris S, Marsteller N, Condon SA, Brown DM. 2014. Early cytokine dysregulation and viral replication are associated with mortality during lethal influenza infection. Viral Immunol 27:214–24.

10. Taubenberger JK, Morens DM. 2008. The pathology of influenza virus infections. Annu Rev Pathol 3:499–522.

11. Kash JC, Taubenberger JK. 2015. The role of viral, host, and secondary bacterial factors in influenza pathogenesis. Am J Pathol 185:1528–36.

12. Magira EE, Chemaly RF, Jiang Y, Tarrand J, Kontoyiannis DP. 2019. Outcomes in Invasive Pulmonary Aspergillosis Infections Complicated by Respiratory Viral Infections in Patients With Hematologic Malignancies: A Case-Control Study. Open Forum Infect Dis 6:ofz247.

13. Rijnders BJA, Schauwvlieghe A, Wauters J. 2020. Influenza-Associated Pulmonary Aspergillosis: A Local or Global Lethal Combination? Clin Infect Dis 71:1764–1767.

14. Schauwvlieghe A, Rijnders BJA, Philips N, Verwijs R, Vanderbeke L, Van Tienen C, Lagrou K, Verweij PE, Van de Veerdonk FL, Gommers D, Spronk P, Bergmans D, Hoedemaekers A, Andrinopoulou ER, van den Berg C, Juffermans NP, Hodiamont CJ, Vonk AG, Depuydt P, Boelens J, Wauters J. 2018. Invasive aspergillosis in patients admitted to the intensive care unit with severe influenza: a retrospective cohort study. Lancet Respir Med 6:782–792.

15. Tobin JM, Nickolich KL, Ramanan K, Pilewski MJ, Lamens KD, Alcorn JF, Robinson KM. 2020. Influenza Suppresses Neutrophil Recruitment to the Lung and Exacerbates Secondary Invasive Pulmonary Aspergillosis. The Journal of Immunology 205(2):480–488.

16. Liu KW, Grau MS, Jones JT, Wang X, Vesely EM, James MR, Gutierrez-Perez C, Cramer RA, Obar JJ. 2022. Postinfluenza Environment Reduces Aspergillus fumigatus Conidium Clearance and Facilitates Invasive Aspergillosis In Vivo. mBio doi:10.1128/mbio.02854-22:e0285422.

17. Sarden N, Sinha S, Potts KG, Pernet E, Hiroki CH, Hassanabad MF, Nguyen AP, Lou Y, Farias R, Winston BW, Bromley A, Snarr BD, Zucoloto AZ, Andonegui G, Muruve DA, McDonald B, Sheppard DC, Mahoney DJ, Divangahi M, Rosin N, Biernaskie J, Yipp BG. 2022. A B1a - natural IgG - neutrophil axis is impaired in viral- and steroid-associated aspergillosis. Science Translational Medicine 14:eabq6682.

18. Guerra ES, Lee CK, Specht CA, Yadav B, Huang H, Akalin A, Huh JR, Mueller C, Levitz SM. 2017. Central Role of IL-23 and IL-17 Producing Eosinophils as Immunomodulatory Effector Cells in Acute Pulmonary Aspergillosis and Allergic Asthma. PLoS Pathog 13:e1006175.

19. Seldeslachts L, Vanderbeke L, Fremau A, Reséndiz-Sharpe A, Jacobs C, Laeveren B, Ostyn T, Naesens L, Brock M, Van De Veerdonk FL. 2021. Early oseltamivir reduces risk for influenza-associated aspergillosis in a double-hit murine model. Virulence 12:2493–2508.

20. Stackowicz J, Jönsson F, Reber LL. 2019. Mouse Models and Tools for the in vivo Study of Neutrophils. Front Immunol 10:3130.

21. Stephens-Romero SD, Mednick AJ, Feldmesser M. 2005. The pathogenesis of fatal outcome in murine pulmonary aspergillosis depends on the neutrophil depletion strategy. Infect Immun 73:114–25.

22. Brunel SF, Bain JM, King J, Heung LJ, Kasahara S, Hohl TM, Warris A. 2017. Live Imaging of Antifungal Activity by Human Primary Neutrophils and Monocytes in Response to A. fumigatus. J Vis Exp 122:e55444.

23. Jhingran A, Mar KB, Kumasaka DK, Knoblaugh SE, Ngo LY, Segal BH, Iwakura Y, Lowell CA, Hamerman JA, Lin X, Hohl TM. 2012. Tracing conidial fate and measuring host cell antifungal activity using a reporter of microbial viability in the lung. Cell Rep 2:1762–73.

24. Sun K, Metzger DW. 2014. Influenza infection suppresses NADPH oxidase-dependent phagocytic bacterial clearance and enhances susceptibility to secondary methicillin-resistant Staphylococcus aureus infection. J Immunol 192:3301–7.

25. Oliveira LVN, Costa MC, Magalhaes TFF, Bastos RW, Santos PC, Carneiro HCS, Ribeiro NQ, Ferreira GF, Ribeiro LS, Goncalves APF, Fagundes CT, Pascoal-Xavier MA, Djordjevic JT, Sorrell TC, Souza DG, Machado AMV, Santos DA. 2017. Influenza A Virus as a Predisposing Factor for Cryptococcosis. Front Cell Infect Microbiol 7:419.

26. Vanderbeke L, Janssen NAF, Bergmans D, Bourgeois M, Buil JB, Debaveye Y, Depuydt P, Feys S, Hermans G, Hoiting O, van der Hoven B, Jacobs C, Lagrou K, Lemiale V, Lormans P, Maertens J, Meersseman P, Mégarbane B, Nseir S, van Oers JAH, Reynders M, Rijnders BJA, Schouten JA, Spriet I, Thevissen K, Thille AW, Van Daele R, van de Veerdonk FL, Verweij PE, Wilmer A, Brüggemann RJM, Wauters J. 2021. Posaconazole for prevention of invasive pulmonary aspergillosis in critically ill influenza patients (POSA-FLU): a randomised, open-label, proof-of-concept trial. Intensive Care Med 47:674–686.

27. van de Veerdonk FL, Gresnigt MS, Romani L, Netea MG, Latgé JP. 2017. Aspergillus fumigatus morphology and dynamic host interactions. Nat Rev Microbiol 15:661–674.

28. Kozel TR, Wilson MA, Farrell TP, Levitz SM. 1989. Activation of C3 and binding to Aspergillus fumigatus conidia and hyphae. Infect Immun 57:3412–7.

29. Steele C, Rapaka RR, Metz A, Pop SM, Williams DL, Gordon S, Kolls JK, Brown GD. 2005. The beta-glucan receptor dectin-1 recognizes specific morphologies of Aspergillus fumigatus. PLoS Pathog 1:e42.

30. Tisoncik JR, Korth MJ, Simmons CP, Farrar J, Martin TR, Katze MG. 2012. Into the eye of the cytokine storm. Microbiol Mol Biol Rev 76:16–32.

31. Fajgenbaum DC, June CH. 2020. Cytokine Storm. N Engl J Med 383:2255–2273.

32. RECOVERY Collaborative Group. 2021. Tocilizumab in patients admitted to hospital with COVID-19 (RECOVERY): a randomised, controlled, open-label, platform trial. Lancet 397:1637–1645.

33. van der Sluijs KF, van Elden LJ, Nijhuis M, Schuurman R, Pater JM, Florquin S, Goldman M, Jansen HM, Lutter R, van der Poll T. 2004. IL-10 is an important mediator of the enhanced susceptibility to pneumococcal pneumonia after influenza infection. J Immunol 172:7603–9.

34. Dewi IM, Cunha C, Jaeger M, Gresnigt MS, Gkountzinopoulou ME, Garishah FM, Duarte-Oliveira C, Campos CF, Vanderbeke L, Sharpe AR. 2021. Neuraminidase and SIGLEC15 modulate the host defense against pulmonary aspergillosis. Cell Reports Medicine 2:100289.

35. Prachanronarong KL, Canale AS, Liu P, Somasundaran M, Hou S, Poh YP, Han T, Zhu Q, Renzette N, Zeldovich KB, Kowalik TF, Kurt-Yilmaz N, Jensen JD, Bolon DNA, Marasco WA, Finberg RW, Schiffer CA, Wang JP. 2019. Mutations in Influenza A Virus Neuraminidase and Hemagglutinin Confer Resistance against a Broadly Neutralizing Hemagglutinin Stem Antibody. J Virol 93:e01639–18.

36. Shepardson KM, Jhingran A, Caffrey A, Obar JJ, Suratt BT, Berwin BL, Hohl TM, Cramer RA. 2014. Myeloid derived hypoxia inducible factor 1-alpha is required for protection against pulmonary Aspergillus fumigatus infection. PLoS Pathog 10:e1004378.

37. Bowman JC, Abruzzo GK, Anderson JW, Flattery AM, Gill CJ, Pikounis VB, Schmatz DM, Liberator PA, Douglas CM. 2001. Quantitative PCR assay to measure Aspergillus fumigatus burden in a murine model of disseminated aspergillosis: demonstration of efficacy of caspofungin acetate. Antimicrob Agents Chemother 45:3474–81.

38. World Health Organization. 2020. WHO information for the molecular detection of influenza viruses. https://cdn.who.int/media/docs/default-source/influenza/molecular-detention-of-influenza-viruses/protocols_influenza_virus_detection_feb_2021.pdf?sfvrsn=df7d268a_5. Accessed February 2021.

39. Ramirez-Ortiz ZG, Prasad A, Griffith JW, Pendergraft WF, 3rd, Cowley GS, Root DE, Tai M, Luster AD, El Khoury J, Hacohen N, Means TK. 2015. The receptor TREML4 amplifies TLR7-mediated signaling during antiviral responses and autoimmunity. Nat Immunol 16:495–504.

